# A novel mechanism of “metal gel-shift” by histidine-rich Ni^2+^-binding Hpn protein from *Helicobacter pylori* strain SS1

**DOI:** 10.1101/081331

**Authors:** Rahul Mahadev Shelake, Yuki Ito, Junya Masumoto, Eugene Hayato Morita, Hidenori Hayashi

**Affiliations:** Proteo-Science Center, Ehime University, Matsuyama, Japan; Laboratory of Molecular Cell Physiology, Faculty of Agriculture, Ehime University, Matsuyama, Japan; Department of Chemistry, Faculty of Science, Josai University, Saitama, Japan

**Keywords:** Histidine-rich protein, Hpn, nickel, anomalous SDS-PAGE migration, gel shifting on SDS-PAGE, MALDI-TOF-MS, protein-metal ion complex

## Abstract

Sodium dodecyl sulphate-polyacrylamide gel electrophoresis (SDS-PAGE) is a universally used method for determining approximate molecular weight (MW) in protein research. Migration of protein that does not correlate with formula MW, termed “gel shifting” appears to be common for histidine-rich proteins but not yet studied in detail. We investigated “gel shifting” in Ni^2+^-binding histidine-rich Hpn protein cloned from *Helicobacter pylori* strain SS1. Our data demonstrate two important factors determining “gel shifting” of Hpn, polyacrylamide-gel concentration and metal binding. Higher polyacrylamide-gel concentrations resulted in faster Hpn migration. Irrespective of polyacrylamide-gel concentration, preserved Hpn-Ni^2+^ complex migrated faster (3-4 kDa) than apo-Hpn, phenomenon termed “metal gel-shift” demonstrating an intimate link between Ni^2+^ binding and “gel shifting”. To examine this discrepancy, eluted samples from corresponding spots on SDS-gel were analyzed by matrix-assisted laser desorption/ionization-time-of-flight mass spectrometry (MALDI-TOF-MS). The MW of all samples was the same (6945.66±0.34 Da) and identical to formula MW with or without added mass of Ni^2+^. MALDI-TOF-MS of Ni^2+^-treated Hpn revealed that monomer bound up to six Ni^2+^ ions non-cooperatively, and equilibrium between protein-metal species was reliant on Ni^2+^ availability. This corroborates with gradually increased heterogeneity of apo-Hpn band followed by compact "metal-gel shift" band on SDS-PAGE. In view of presented data metal-binding and “metal-gel shift” models are discussed.

## Introduction

Cysteine-rich (-SH group) metal-binding proteins, for example metallothioneins, are known to bind and sequester multiple metal ions in prokaryotes and eukaryotes [1], but understanding of histidine-rich (imidazole group) metal-binding proteins is still limited. Proteins with histidine-rich repeats are universally present in the proteomes of both prokaryotes and eukaryotes, including human, and form unique single-residue-repeat motifs [1,2]. *Helicobacter pylori*, a human pathogen responsible for severe gastric diseases including intestinal ulcers and adenocarcinoma, requires two key Ni-containing enzymes (urease and hydrogenase) for survival in acidic gastrointestinal conditions [3]. Thus, *H. pylori* produce unique Ni^2+^-binding histidine-rich proteins required in the maturation of urease and hydrogenase.

One of the first candidates involved in Ni^2+^ homeostasis in *H. pylori* was isolated and named Hpn (<u>*H*</u>*elicobacter* <u>*p*</u>*ylori* protein binding to <u>n</u>ickel) [4]. Hpn contains 28 histidine residues (46.7%) with two stretches of repeated histidines at positions 11-17 and 28-33 (**Fig 1A**). It also contains two short repeating motifs (EEGCC) in the internal part positioned at 38-42 and 51-55. The relative affinity of Hpn towards divalent metal ions was found to be different under *in vivo* and *in vitro* conditions indicating the complex nature of the protein [4– 7]. Initial Hpn mutation studies in *H. pylori* showed that the mutant strain was more sensitive to Ni^2+^ than a wild-type strain [4,8,9]. Inductively coupled plasma-mass spectrometry and equilibrium dialysis studies revealed that average five Ni^2+^ ions (5.1±0.2) bind to Hpn in a pH-dependant manner and forms a range of multimeric complexes (>500, 136, 55, 34, 26, 20, 14 and 7 kDa) in solution that exists in equilibrium depending on buffer content [5,6]. The pH titration and competition experiments using EDTA confirmed that the metal-binding to Hpn is a reversible process [5–7]. Even though some amino-acid residues vital for metal-binding were identified [10–12], distribution of metal-binding sites and equilibrium between protein-metal species in Hpn is not yet known. Hpn may act in Ni^2+^ storage as a ‘reservoir’ or in channelizing Ni^2+^ to other proteins [5,6,13]. Purified Hpn can form amyloid-like fibers *in vitro* [14] but such fibers yet to be established *in vivo* in *H. pylori*. A recent study described Hpn interaction with Hpn-2 (also known as Hpn-like or Hpn-l, another atypical histidine-rich protein with MW 8.07 kDa [15]) mediate urease activity [13]. Thus, considering several complex chemical interactions, Hpn can be employed as an ideal protein to investigate metallochemistry and physicochemical aspects of histidine-rich metal-binding proteins.

**Fig 1.**
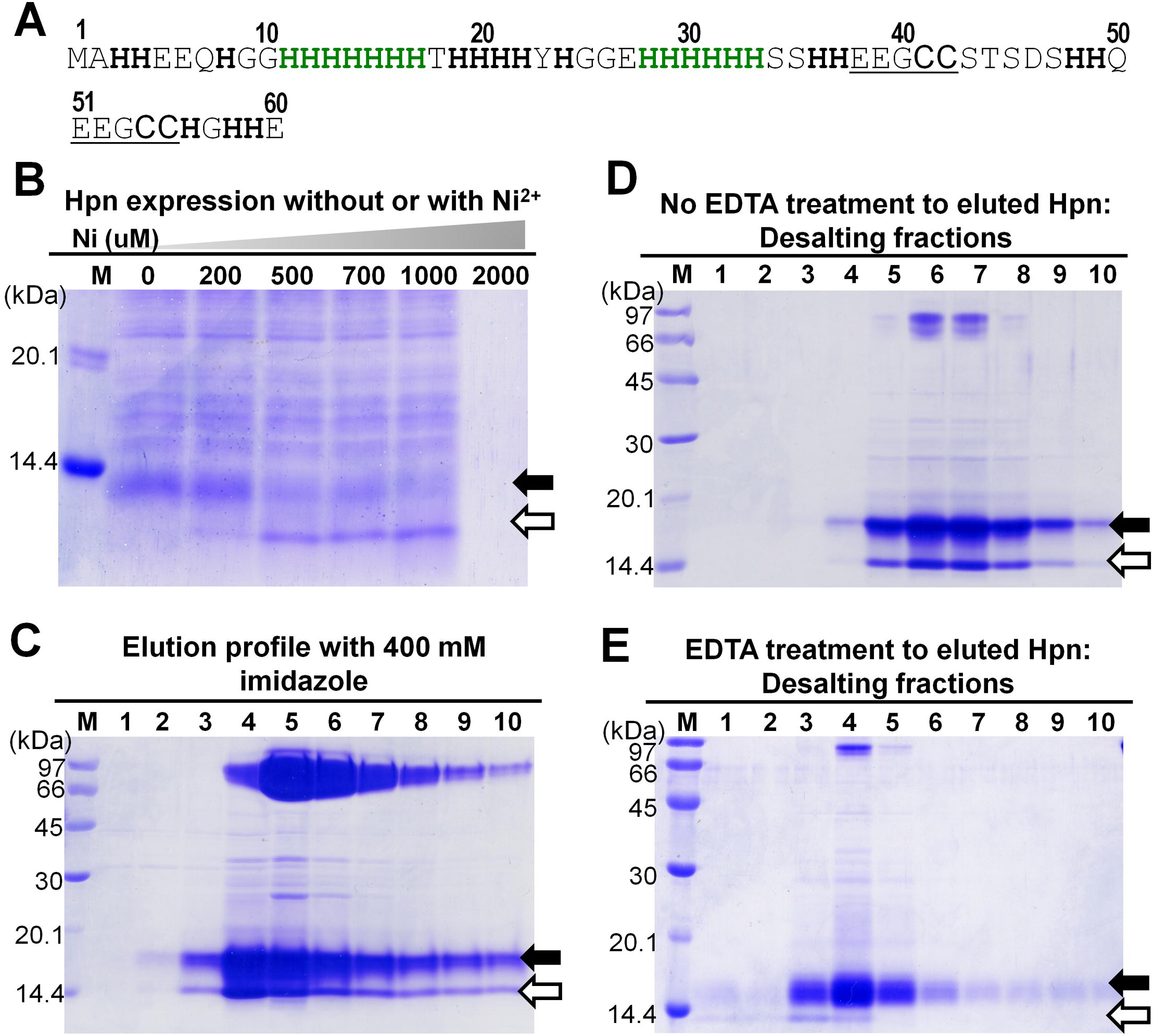
Amino acid sequence, overexpression and purification of recombinant Hpn. Lane M, LMW protein marker standards (GE Healthcare); black arrows depicting apo-Hpn and white arrows showing probable Ni^2+^-bound Hpn protein in all panels. 1. Amino acid sequence of Hpn. Histidine residues are highlighted in bold. Stretches of six and seven histidines are highlighted in green and pentapetide repeats (EEGCC) are underlined. 2. SDS-PAGE of Hpn expression with or without Ni^2+^ added in the culture (polyacrylamide-gel 20%). Pellets of 60 µl bacterial cultures were dissolved in 60 µl of 1X Laemmli buffer and boiled for 3 min at 100ºC. Final volume of 15 µl loaded in each lane. C, D and E. Elution profile of purified Hpn checked by loading protein fractions on SDS-PAGE (polyacrylamide-gel 15%). Lanes 1 to 10, fractions of purified protein eluted with 400 mM imidazole (C). Elution profiles of desalted fractions of Hpn without EDTA treatment (D) and with EDTA-treatment (E) were analyzed. Equal volume of 2X Laemmli buffer was added to each eluted fraction and then boiled for 3 min at 100ºC. Total 10 µl applied in each lane in C, D and E.

Even though sodium dodecyl sulfate-polyacrylamide gel electrophoresis (SDS-PAGE) is the most commonly used method for the determination of approximate MW of proteins; unusual electrophoretic behavior in SDS-PAGE has been reported for Hpn, making its identification problematic [5]. This phenomenon of unpredictable migration rate on SDS-PAGE against actual formula MW of protein is termed “gel shift” (not to be confused with the electrophoretic mobility shift assay for protein-DNA interactions). The “gel shifting” has also been reported for several other histidine-rich proteins including Hpn [5], UreE [16], HypB [17–19], SlyD [20,21], Hpn-like [9], and CooC [22] **(Table 1)** as well as different helical membrane proteins [23–25] including some isobaric peptides (peptides with the same MW) from various organisms [26]. HspA from *Helicobacter*, which has a unique C-terminus containing a histidine-rich region, also exhibited a higher MW of 15.5 kDa than the expected 13 kDa [27] indicating that the “gel shift” behavior might be attributable to histidine-rich region. However, the molecular mechanism is yet to be clarified. Six possible mechanisms responsible for “gel shifting” of a protein have been hypothesized in previous reports: (a) divergent higher order (secondary or tertiary) structure; (b) difference in Stokes-Einstein (hydrodynamic) radius of the protein-surfactant complex; (c) variation in the intrinsic net charge of the protein; (d) number of bound SDS molecules; (e) post-translational modifications; or (f) binding of cofactors such as metal ions to the protein [23–25,28–30]. Nevertheless, polyacrylamide-gel concentration can dictate magnitude and direction of some proteins in SDS-gel [31].

**Table 1.** List of histidine-rich Ni^2+^-binding proteins showing gel shifting anomaly.

| Protein | Organism | Function | Histidine content | Ni ions per monomer | Formula MW (kDa) | Apparent MW (kDa) | Reference |
| --- | --- | --- | --- | --- | --- | --- | --- |
| SlyD | *Escherichia coli* | Ni carrier | 10.2% | 3 | 20.8 | 25 | [20,21] |
| CooC | *Rhodospirillum rubrum* | Nickel accessory protein for maturation of CODH | 3% | 1 | 27.8 | 61, 29 | [22] |
| HypB | *Rhizobium leguminosarum* | GTP-dependent Ni insertase | 9.0% | 4 | 32.5 | 39 | [18,19] |
| HypB | *Archeoglobus fulgidus* | GTP-dependent Ni insertase | 2.7% | 0.5 | 24.7 | 41 (dimer) | [17] |
| UreE | *Klebsiella aerogenes* | Accessory protein in urease maturation | 9.5% | 3 | 17.5 | 35 (dimer) | [16] |
| Hpn like | *Helicobacter pylori* | Ni storage | 25% | 2 | 9 | 47, 18 | [9] |
| Hpn | *Helicobacter pylori* | Ni storage | 46.7% | 5-6 | 7 | >670, 500, 230, 136, 20, 14, 7 | Present study and [5] |

Recently, attempts have been made to investigate these molecular mechanisms using segments of various membrane proteins, and the findings indicated that there might be no single universal mechanism that accounts for the anomalous migration of all proteins. In fact, each protein or class of proteins may have unique chemistry responsible for “gel shifting” behavior. It is an interesting question that whether “gel shifting” in Hpn is linked to the displacement of protein-protein, protein-metal, and/or protein-SDS interactions, but these need further investigation.

We cloned the *hpn* gene from the Sydney Strain 1 (SS1) of *H. pylori*, a standardized mouse model [32]. Our preliminary data demonstrated the altered migratory position for Hpn on SDS-PAGE when Hpn was expressed in Ni^2+^-supplied medium suggesting role of Ni^2+^ in altered migration. We find that the migration rate of Ni^2+^-treated Hpn on SDS-PAGE was altered because of preserved protein-metal complexes. Preservation of intact protein-metal bond in non-denaturing conditions is commonly reported [33,34]. SDS-PAGE is primarily designed to separate denatured proteins by disrupting non-covalent complexes. However, in SDS-treated samples, protein under study and proteins used as MW standards attain equivalent shape not by an absolute unfolding but indeed it is achieved by the SDS aggregation at hydrophobic sites [23]. Thus, protein denaturation occurs in ‘‘reconstructive’’ manner and it forms a mixture of SDS micelle wrapped around α-helices separated by linkers (necklace and bead model) or hydrophobic regions of protein wrapped around SDS micelles (decorated micelle model), both reviewed in [35]. Thus, some SDS-resistant protein-protein interactions (**supporting information, S1 Table in S1 file**) and protein-metal complexes (**supporting information, S2 Table in S1 file**) can also preserve in electrophoretic separation upon SDS-PAGE provided interaction is stronger [34]. Also, many studies reported that the “reconstructive” denaturation can retain some protein-metal complexes on SDS-PAGE but altered migration is not generally observed except for some Ca^2+^-binding proteins such as calmodulin isoforms [36] and Ca^2+^-dependent protein kinases (CDPKs) [37–40]. Overall, the above observations suggest that there may be an apparent link between metal-binding to Hpn and “gel shifting” pattern. Thus, the anomalous migration of Hpn should be explored in several conditions pertaining to metal binding and its effect on migration rate in SDS-PAGE. Also, MW of intact Hpn alongside protein-metal interaction studies by MS could facilitate better understanding of Hpn metallochemistry in detail.

In the present study, an attempt was made to investigate various physicochemical properties of Hpn and not the *in vivo* function of Hpn in *H. pylori* itself. This work was focused on heterologous expression of Hpn in *E. coli*, average molecular mass of purified Hpn, “gel shifting” anomaly with or without Ni^2+^ in different polyacrylamide-gel concentrations and application of MALDI-TOF-MS for studying non-covalent Hpn-Ni^2+^ complexes. To establish a reliable technique to determine MW associated with “gel shifting” anomaly without ambiguity, MW of intact recombinant Hpn was determined by MALDI-TOF-MS using internal protein standards and found to be essentially identical (6945.66±0.34) to the formula MW. Ni^2+^-treated Hpn migrated more rapidly than untreated Hpn showing differences of 3-4 kDa. This significant metal-triggered shift in electrophoretic gel mobility of Hpn is reported for the first time to the best of our knowledge and termed as a “metal gel-shift”. Migration speed of Hpn in both the forms was altered depending on polyacrylamide-gel concentrations. The method for sample and matrix preparation standardized in this study preserved appropriate intermolecular interactions and hence boosts the usefulness of MALDI for the study of non-covalent protein-metal ion complexes.

## Materials and Methods

### *H. pylori* strain and growth conditions

All the chemicals and reagents used were of analytical grade or higher. The mouse-adapted strain of *Helicobacter pylori* Sydney strain (*H. pylori* SS1) was used in this study [32]. *H. pylori* SS1 was grown on Tripticase Soy agar with 5% sheep blood (TSA II) (Becton Dickinson, Franklin Lakes, NJ) for three days at 35ºC in 12% carbon dioxide condition. A single colony was isolated and sub-cultured on TSA II agar at 35ºC in 12% carbon dioxide condition.

### Purification of DNA from *H. pylori* strain SS1

The genomic DNA from strain SS1 was extracted based on phenol-chloroform method as described previously [32] with minor modifications. Briefly, bacteria were harvested from TSA II agar and suspended in 3 ml of Tris-buffered saline (TBS). After washing with TBS once by centrifugation at 3,800 g for 5 min at 4ºC, 5 x 10^8^ colony forming unit (CFU) of bacteria were re-suspended in 500 μl of lysis buffer (50 mM Tris-HCl, pH 8.0; 100 mM EDTA, pH 8.0; 1 % SDS, 100 mM NaCl) containing 0.2 mg/ml proteinase K (Amaresco, Solon, OH) and incubated at 37ºC for 12 h. Subsequently, UltraPure™ buffer-saturated phenol (Invitrogen, Carlsbad, CA) was added, and the mixture was gently rotated for 15 min. After centrifugation, the aqueous phase was transferred to a tube containing chloroform/isoamyl alcohol (24:1, Sigma-Aldrich, St. Louis, MO) and agitated gently for 10 min. The aqueous phase was collected to a new tube by centrifugation. The DNA was precipitated by addition of isopropyl alcohol and 0.3 M sodium acetate, and then the DNA pellet was rinsed with 70% ethanol. After centrifugation, pellet was air-dried and dissolved in Tris-EDTA buffer (pH 8.0).

### Cloning and nucleotide sequencing

First, the Hpn encoding region of 1.17 kb was amplified instead of the *hpn* gene only to avoid undesired mutations in the nucleotide sequence of *hpn* (primers 5'-AGTCCATATGCCTTACACGCCGTAGATGACAAAACGCGC-3' and 5'-GACTGGATCCGGCTCGCTCTCATCTATAGCGTGGCTAAG-3'). The DNA fragment was cloned into a pBluescript II KS (+) vector (Takara, Japan), and nucleotide sequencing was performed using a BigDye Terminator v3.1 Cycle Sequencing Kit and an ABI PRISM 310 Genetic Analyzer (Applied Biosystems). The *hpn* gene was amplified using the cloned 1.17 kb region as the PCR template (primers 5'-GACTCATATGGCACACCATGAAGAACAACAC-3' and 5'-GACTGGATCCTTATTACTCGTGATGCCCGTGGC-3') and then cloned into pBluscript (Takara, Japan) and pET21b (Novagen, Darmstadt, Germany) vectors. *E. coli* JM 109 and BL21 (DE3) cells purchased from Takara (Japan) were used for the propagation and protein expression with a constitutive (RuBisCo promoter from *Synechococcus* sp. PCC7002) [41]; and isopropyl β-D-1-thiogalactopyranoside (IPTG)-inducible T7 promoter, respectively.

### Expression and purification of recombinant Hpn

Recombinant Hpn was purified from harvested cells of 200 ml culture. Harvested cells were suspended in lysis buffer (50 mM Tris-HCl, pH 7.5, and 500 mM NaCl) and purification was done as described previously for His-tagged proteins with some modifications [42]. After sonication (TOMY Ultrasonic Disruptor, duty: 50, output: 4, time: 4 min x 6), supernatant fractions were filtered using Millex^®^GV filter units of 0.22-µm-pore-size and applied onto a 1-ml HiTrap chelating column (GE Healthcare) that had been equilibrated with start buffer containing 20 mM imidazole, 50 mM Tris-HCl, and 500 mM NaCl. The column was washed with buffer (50 mM Tris, pH 7.5, 50 mM NaCl, and 40 mM imidazole), and then Hpn was eluted using start buffer with 400 mM instead of 20 mM imidazole. Eluted fractions were analyzed for purity using SDS-PAGE (15%), and the purest fractions were applied onto a 5-ml HiTrap desalting column (GE Healthcare) that had been equilibrated with desalting buffer (20 mM Hepes-KOH, pH 7.4, 100 mM NaCl, and 20% glycerol), and purified Hpn was stored at −80ºC until use. The concentration and quality of purified Hpn was measured using bovine serum albumin (BSA) as the standard in a BCA assay (Bio-Rad) in accordance with the manufacturer’s instructions and SDS-PAGE (polyacrylamide 15%), respectively.

### SDS-PAGE and non-denaturing native-PAGE (native-PAGE)

All purified protein fractions were analyzed by SDS-PAGE in accordance with instructions mentioned previously [43]. A vertical electrophoresis system of mini-slab size from Atto Co. (Japan) was used for the separation of protein samples with appropriate concentrations of polyacrylamide. A LMW-SDS Marker Kit (GE Healthcare) and marker proteins kit from Nacalai Tesque, Inc. (Japan) were used as protein standards in SDS-PAGE. All protein samples were prepared in Laemmli buffer (50 mM Tris-HCl pH 6.8, 10% glycerol, 2% SDS, 7% β-mercaptoethanol and 0.001% bromophenol blue) and then stored at −20ºC before use. The electrophoresis was done till bromophenol blue dye reached to the bottom in all gels of different concentrations (under similar experimental conditions). This optimized method was referred from previous work [31] showing separation of 11 trans-membrane mimetic peptides, translating into MWs of 3.5-41 kDa on 11-18% of polyacrylamide-gel. Resolved proteins were stained with Coomassie Brilliant Blue (CBB) R-250 solution containing 10% acetic acid with 25% methanol.

Fractions of purified Hpn were used for blue native PAGE analysis without boiling or reduction in sample buffer (50 mM Tris with pH 6.8 containing 0.01% bromophenol blue and 10% glycerol). The running buffer was prepared without detergent and consisted of 25 mM Tris, 192 mM glycine, pH 8.0. Stacking gel (4%) and separation gel (10%) were prepared adding suitable buffers, glycerol, ammonium persulfate (APS), and TEMED. Electrophoresis was performed at a constant current of 15 mA and voltage of 160 V and further CBB stained.

### Western blotting analysis

The Hpn protein (25 µM) treated with indicated mol equivalent of EDTA or Ni^2+^ resolved on SDS-PAGE (20%) and then electrophoretically blotted onto polyvinylidene fluoride (PVDF) membrane (Amersham Hybond-P, GE Healthcare, code: RPN303F) in transfer buffer containing no methanol. Blotting was performed at 0.8 mA current for every cm^2^ area of gel. Proteins were blocked on PVDF membranes in a solution of 5% skim milk for 1 h after washing in PBS-T buffer (phosphate-buffered saline buffer with 0.05% Tween-20, pH 8). The PVDF membrane was further incubated with C-terminal specific-anti 6×histidine monoclonal antibody (9F2) (Wako Japan, product code: 010-21861 diluted to 1:5000 in PBS-T) for 1 h. After three washes of 15:5:5 min in PBS-T, PVDF membrane was then incubated with horseradish peroxidase-conjugated anti-mouse IgG (GE Healthcare, product code: NA931VS, diluted to 1:5000 in PBS-T) for 1 h. The membrane was further washed again with PBS-T as mentioned earlier and Amersham ECL non-radioactive western blotting detection reagents (GE Healthcare) were used for visualizing protein bands in accordance with the manufacturer’s instructions.

### Enzyme-linked immunosorbent assay (ELISA) analysis

Green fluorescent protein with artificial His.tag (*gfp-His*_*6*_) and GFP fused with Hpn (*gfp-Hpn*) were cloned in pET21b. IPTG-induced over-expression of both the proteins was done with or without Ni^2+^ added in the culture (**Supporting information, S1 Fig in S1 file**). Pellets of 60 µl bacterial cultures were dissolved in 60 µl sample buffer and incubated for 3 min at 100ºC. Final volume of 15 µl loaded in each lane for SDS-PAGE. ELISA experiment was performed using previously described protocol with some modifications [44]. Different buffers were prepared before starting ELISA experiment (components of buffers summarized in Table 2).

**Table 2.** Components of different buffers used in ELISA.

| Buffer type | Components |
| --- | --- |
| Coating buffer | Phosphate buffer saline, [1.16 g Na<sub>2</sub>HPO<sub>4</sub>, 0.1 g KCl, 0.1 g K<sub>3</sub>PO<sub>4</sub>, 4.0 g NaCl (500 ml distilled water) pH 7.4] |
| Dilution buffer | 0.1% BSA with 0.05% Tween-20 in PBS |
| Blocking buffer | 1% BSA with 0.05% Tween-20 in PBS |
| Washing buffer | 0.05% Tween-20 in PBS |
| Stop buffer | 5N Sodium Hydroxide in distilled water |

Flat-bottom 96-well ELISA plates (untreated 96-well microplates from Falcon) were used for coating. Concentration of each protein (Hpn, GFP-Hpn and GFP-His_6_) was adjusted to 1 µg by dilution with coating buffer to the final volume of 50 µl. Plates were incubated at 4ºC for overnight. Next day, solution was thrown away and 200 µl blocking buffer into each well was added. Then, plate was incubated at 37ºC for 1hr. After incubation, solution was discarded and plate was washed three times by washing buffer. His.Tag® antibody was diluted to standardized concentration (1:500) with dilution buffer [C-terminal specific-anti 6xhistidine monoclonal antibody (9F2) (Wako Japan, product code: 010-21861)] and plate incubated at 37ºC for 1hr after adding 100 μl in each well. After three washes with wash buffer, plate was incubated with horseradish peroxidase-conjugated anti-mouse IgG (GE Healthcare, product code: NA931VS, diluted to 1:1000 in wash buffer) for 1 hr at 37ºC. After similar washing, the substrate ABTS [2,2'-Azinobis(3-ethylbenzothiazoline-6-sulfonic Acid Ammonium Salt) from Wako Japan] dissolved in 0.1 M citrate buffer and hydrogen peroxide (0.03%) was added. Then the plate was incubated for 20 minutes at room temperature. Reactions were stopped by adding stop buffer (100 µl) in each well. The absorbance at 415 nm was measured using a Spectramax M3 microplate reader (Molecular Devices Co., Sunnyvale, CA). Values obtained (absorbance at 415 nm) for Ni^2+^-treated samples (from average of at least three replications) were normalized against untreated samples and plotted in graph.

### Hpn interaction with Ni^2+^ ions

For SDS-PAGE and blue native-PAGE analysis, apo-Hpn (25 µM) was prepared as mentioned above and then treated with the indicated amount of Ni^2+^ by adding NiSO_4_ solution or EDTA (with mol equivalent ratio of 1:6) and incubated for a minimum of 1 h at room temperature. Suitable PAGE buffer was added to the above mixture for loading either on SDS-PAGE (Laemmli buffer) or blue native PAGE (sample buffer used, 50 mM Tris, pH 6.8 with 10% glycerol and 0.01% bromophenol blue). Equal volume of 2X loading buffer was added to protein samples and then applied to native-PAGE directly. For SDS-PAGE, further processing (heat denaturation) done otherwise mentioned in respective figures. Ni^2+^ binding to Hpn protein was investigated using MALDI-TOF-MS by mixing Ni^2+^ or EDTA-treated Hpn with equal amount of matrix solution.

The gel slices of EDTA or Ni^2+^-treated protein bands resolved on SDS-gel were digested in nitric acid (Nacalai Tesque, Japan) and heated at 80ºC for 10 min, once cooled nitric acid finally diluted to 2%. The Ni^2+^ content was analyzed by ICP-optical emission spectrometry (ICP-OES) (Optima 8300 ICP-OES Spectrometer, PerkinElmer Inc, USA). The standard curve was plotted using a minimum of five standards (ranging from 50 to 1000 ppm) with a blank of 2% nitric acid.

### MALDI-TOF-MS

Protein bands resolved on SDS-PAGE were eluted using the protocol described previously [45] with or without CBB staining and then aliquots were used in mass spectrometry. Purified Hpn and elution fractions were directly mixed with an equal volume of matrix solution prepared with different recipes. Acidic and non-acidic matrices were investigated for detecting intact protein-metal ion complexes. Acidic matrix was prepared by mixing 100% acetonitrile (ACN), 0.1% trifluoroacetic acid (TFA), and distilled water (v:v, 50:10:40). Another mild acidic matrix was prepared with the recipe except for more diluted 0.01% TFA. Two different non-acidic matrices were analyzed: 1) sinapinic acid in 100% ACN and distilled water (v:v, 1:1) without TFA [46]; and 2) 4-hydroxy-α-cyanocinnamic acid powder in a saturated solution of ethanol and 1 M ammonium acetate (v:v, 1:1) [47]. MALDI-TOF-MS analysis was done with a Voyager-DE PRO MALDI-TOF-MS (Applied Biosystems). The instrument was calibrated externally with a Sigma Protein MALDI-MS calibration kit. The positive linear mode was set in the instrument to acquire mass spectra using a nitrogen laser (337 nm). An average of a minimum of 150 laser shots was used to accumulate a single spectrum with an accelerating voltage of 25,000 V and extraction delay time of 400 ns. Internal calibration with protein standards (insulin and apomyoglobin), smoothing and baseline correction of the mass spectra was performed and analyzed by using Data Explorer software (Applied Biosystems, MA).

### Effect of Ni^2+^ on growth of *E. coli* expressing the *hpn* gene

*E. coli* cultures, both with pBluscript only and pBluscript-*hpn* plasmid inoculated from individual colonies were grown for overnight in Luria-Bertani (LB) medium containing 100 μg/ml ampicillin at 37ºC. Optical density (OD) of the grown cultures was measured at 600 nm and normalized to 1, of which 100 µl was inoculated to 5 ml of fresh LB medium (1:50 dilution) containing 100 μg/ml ampicillin, and NiSO_4_ was added where applicable (0, 500, 1000, 1200 μM). Cultures were grown for 4 h at 165 rpm at 37ºC, OD values were measured using U-1800 spectrophotometer (Hitachi, Japan) at 600 nm. Obtained values were normalized against control (culture grown without Ni^2+^ stress) and data from three different replicates was summarized and used to plot the growth curve.

M9 medium is a minimal defined culture medium prepared as described previously [48]. Overnight grown cultures (as above, 100 μl) were inoculated into 5 ml of fresh M9 medium containing 0, 50, and 100 μM NiSO_4_ and 100 μg/ml ampicillin. These cultures were incubated for 24 h at 165 rpm at 37ºC in a shaker and the growth curve was plotted (as described above).

### Ni^2+^ accumulation in *E. coli* expressing the *hpn* gene

Single colonies of transformed *E. coli* JM109 were inoculated into 2 ml of fresh medium (LB or M9) supplemented with appropriate concentrations of Ni^2+^. Overnight grown cultures (after normalizing OD to 1) were inoculated into 5 ml of fresh medium and grown for 24 h at 37ºC with and without Ni^2+^. Sampling (2 ml culture) was done at 24 h (LB and M9), respectively. Cells were centrifuged at 15000 rpm for 1 min and bacterial pellet was washed with GET buffer (50 mM glucose, 25 mM Tris-HCl, 10 mM EDTA) followed by drying at 80ºC for a minimum of 3 h. The dried pellet was digested in nitric acid (Nacalai Tesque, Japan) and heated at 80ºC for 10 min, once cooled nitric acid finally diluted to 2%. The Ni^2+^ content incorporated inside the cell and total uptake were analyzed by ICP-OES.

## Results

### Overexpression and purification of Hpn

Four changes were observed at the nucleotide level in the *hpn* ORF of strain SS1 compared with strain 26695, but it had no change at the amino-acid level (**Supporting information, S2 Fig in S1 file**). Hpn expression in *E. coli* culture was standardized in LB medium. The molecular mass of Hpn expressed with and without Ni^2+^ was investigated by SDS-PAGE with a crude cell extract and found to be different (**Fig 1B**). The SDS-PAGE pattern of imidazole-eluted Hpn showed three major protein bands of approximately 14 kDa, 18 kDa, and 70 kDa on 15% polyacrylamide-gel (**Fig 1C**). The SDS-resistant oligomeric complex (~70 kDa) was highly stable and unaffected by reducing agents such as β-mercaptoethanol or boiling (100ºC for 3 min). After desalting, this ~70 kDa band was observed but at very low concentration, thereby suggesting the role of imidazole in inter-conversion of Hpn multimeric forms (**Fig 1D**). The other two protein bands of ~14 kDa and ~18 kDa were observed corresponding to the pattern of the crude extract with and without Ni^2+^ respectively. This indicates that a trace of Ni^2+^ was present in purified Hpn incorporated during the purification process. Thus, the purified Hpn was used after removing trace amounts of Ni^2+^ by treating with EDTA during buffer exchange with desalting columns (**Fig 1E**).

It has been shown that Hpn forms a range of multimeric complexes depending on buffer composition and treatment of DTT, imidazole and Ni^2+^ estimated by gel-filtration chromatography [5]. Consistent with earlier report, Hpn protein (25 μM) treated with either EDTA or Ni^2+^ (mol equivalent ratio of 1:6 independently) migrated as a range of multimeric species on 10% native-PAGE gel (**Supporting information, S3Fig in S1 file**).

### Confirmation of recombinant Hpn by Western blotting

The post-translational removal of N-terminal methionine (149.21 Da) in wild-type (in *Helicobacter pylori*) and recombinant Hpn (in *E. coli*) is reported [4,5]. Therefore, the monoisotopic and average molecular mass (abbreviated as *M_MONO_* and *M*_*av*_) of Hpn without N-terminal methionine were estimated using PAWS software (http://www.proteometrics.com) as 6941.288 Da and 6946.01 Da, respectively. Hpn appeared to migrate faster in the sample from cells cultured with Ni^2+^ compared to that without Ni^2+^ on 15% (**Fig 1C-E**) or 20% SDS-PAGE (**Fig 1B**), but neither of the conditions showed the migratory position for an expected MW i.e. ~7 kDa.

To facilitate the identification of Hpn protein (untagged) precisely, western blotting was performed using a His.Tag^®^ monoclonal antibody assuming that the His.Tag^®^ antibody would bind to Hpn at the seven and six residue histidine repeats [5]. Recombinant Hpn (untagged) confirmed a thin band in western blot (**Fig 2A**, left panel) corresponding to the migration position of apo-Hpn but Ni^2+^-treated Hpn was barely visible. CBB staining of same PVDF membrane showed both the protein bands but small amount of Ni^2+^-treated Hpn remain bound to membrane (**Fig 2A**, right panel). Possible explanation is that untagged Hpn, specifically metalated form may have weaker binding affinity to membrane probably due to its atypical chemical nature.

**Fig 2.**
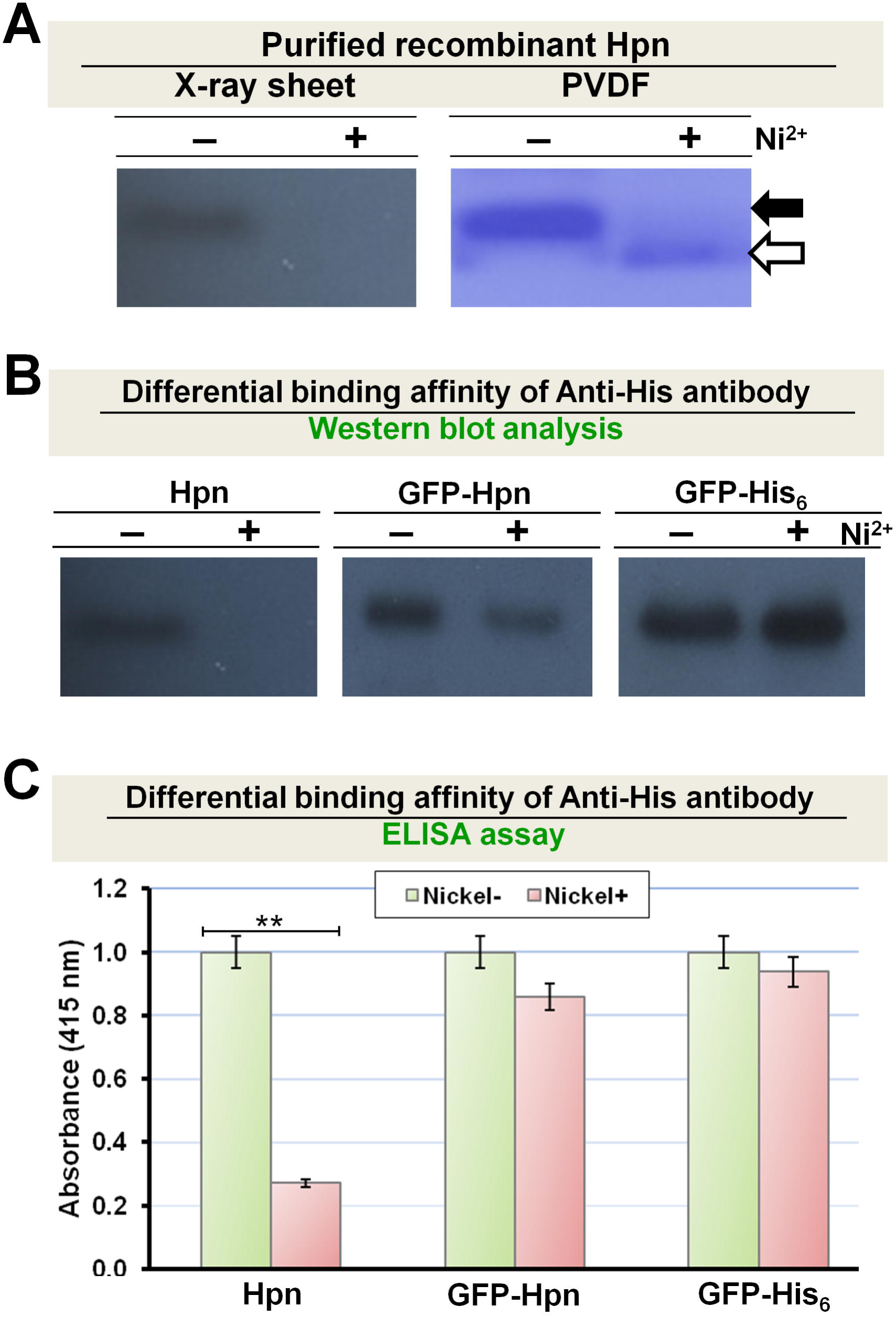
Western blot and ELISA analysis of Hpn, GFP-His_6_ and GFP-Hpn. A. Western blot of Hpn was done using His.Tag^®^ monoclonal antibody. In left panel, X-ray sheet showing ECL detection result of recombinant Hpn (with and without Ni^2+^) and right panel showing same CBB-stained-PVDF membrane used for ECL detection. B. Western blot of untagged Hpn, GFP-Hpn and GFP-His_6_ on PVDF membrane either treated with or without Ni^2+^ solution (1mM of NiSO_4_). C. ELISA of denatured untagged Hpn, GFP-Hpn and GFP-His_6_ Equal amount of protein (1 µg) was coated on ELISA plate. His.Tag® antibody was diluted to 1:500 for detection. Values obtained (absorbance at 415 nm) for Ni^2+^-treated samples (from average of at least three replications) were normalized against untreated samples and plotted in graph. Paired t-test was performed to compare the metal ion effect, ** indicate p<0.01.

### Metal-binding to Hpn changes Hpn-antibody interaction

Interaction of His.Tag® antibody with Hpn may not necessarily show similar results for other protein having artificial His.Tag or Hpn conjugated with another protein owing to differential co-ordination geometry of metal-binding and its chemical surrounding. We investigated this possibility using GFP-His_6_ and GFP-Hpn (**Fig 2B**). GFP does not interact with His.Tag® antibody by its own. GFP is comparatively large protein (26.7 kDa) but still positional shift was observed in case of GFP-Hpn expressed in LB medium supplied with Ni^2+^. Western blot data of GFP-His_6_ and GFP-Hpn showed almost equal intensity signals in both the cases i.e. with or without Ni^2+^.

Recognition site for His.Tag® antibody is His_6_ peptide attached at C-terminal of a recombinant protein and it is a linear epitope. Hence, conformational change upon Ni^2+^-binding to Hpn may lead to altered binding of His.Tag® antibody. This was further examined by ELISA using His.Tag® antibody (**Fig 2C**). The SDS-treated and non-treated Hpn protein shown similar results which is compatible with previous observations for several other proteins [49,50]. The relative detection sensitivity of Ni^2+^-treated protein with His.Tag® antibody in ELISA give order of untagged Hpn < GFP-Hpn < GFP-His_6_. Hence, the variability that we observed in detection of apo- and metalated-Hpn on western blots may have resulted not only from membrane-binding efficiency but also from differential exposure of His-rich region upon metal-binding.

These data signify that the metal-binding to Hpn causes altered binding of His.Tag® antibody, possibly due to change in protein confirmation.

### Determination of MW by MALDI-TOF-MS

To quantify the reliable molecular mass of untagged Hpn, MALDI-TOF-MS was used for purified apo-Hpn with an acidic matrix in linear positive mode. As shown in **Fig 3A**, integral signals of four heteromeric species of Hpn (M^+^, 2M^+^, 3M^+^, and 4M^+^) were observed with a major peak of singly charged Hpn[M^+^]. Although a tendency of apo-Hpn to form multimeric complexes in native and reduced states was confirmed by native-PAGE and SDS-PAGE (with 400 mM imidazole-buffer), respectively, the appearance of a major peak of monomer and low intensity peaks of heteromeric species in MALDI-TOF-MS analysis is of a great advantage in determining the molecular mass accurately.

**Fig 3.**
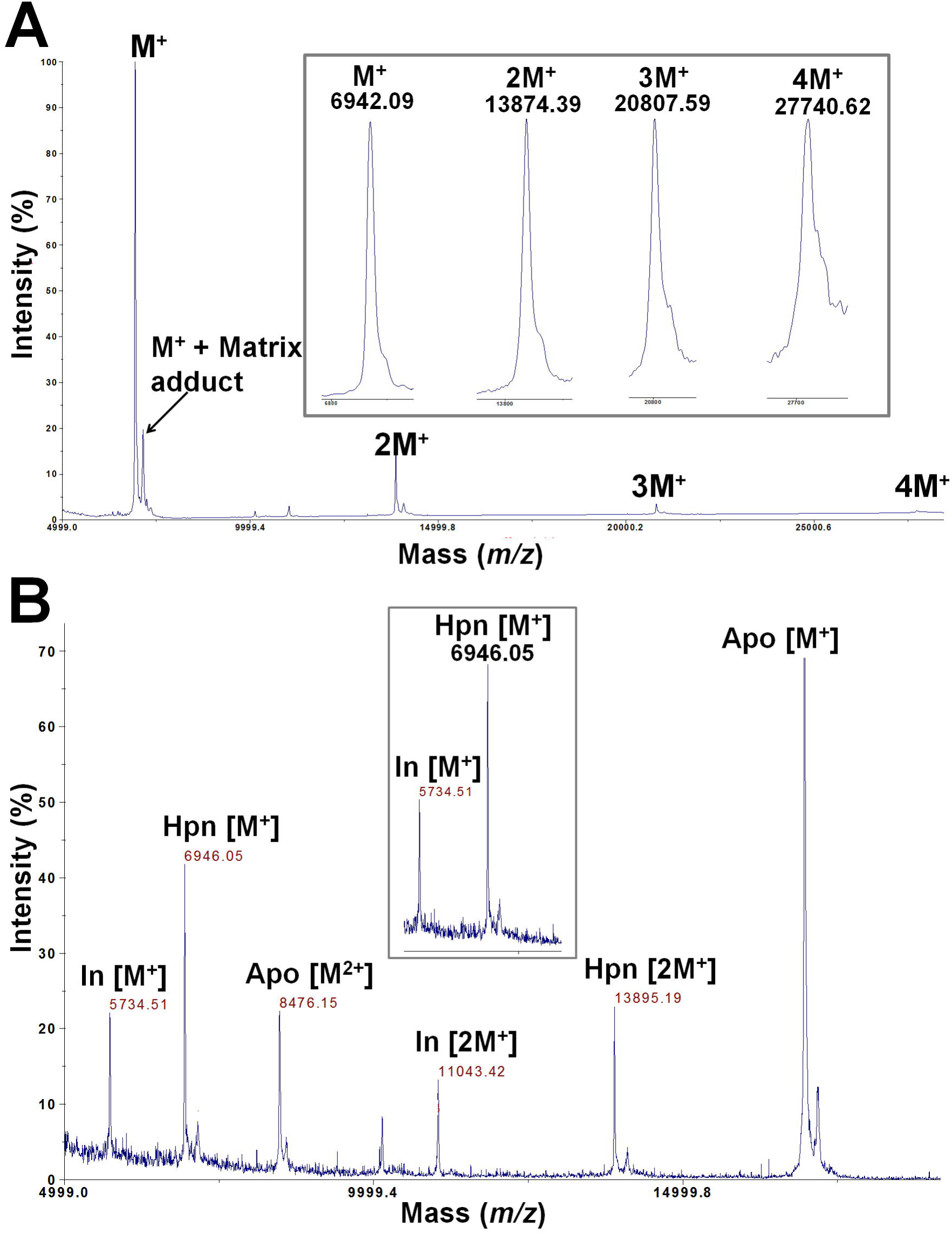
Molecular mass analysis of Hpn with MALDI-TOF-MS. P. Sharp peak of monomeric Hpn together with neighboring small peak of matrix adduct, and three oligomeric species with lower intensity was observed in Hpn spectrum. The monomeric and three oligomeric species: M^+^, 2M^+^, 3M^+^, and 4M^+^ with masses of *m/z* 6942.09, 13874.39, 20807.59, 27740.62 respectively are shown in inset. Q. Molecular mass of Hpn was measured (*m/z* 6946.05) using two different internal standards (insulin and apomyoglobin), which showed peaks for protonated and doubly charged species. Average MW of recombinant Hpn (without methionine) determined (6945.66±0.34) is showing almost negligible difference (0.35) compared to theoretical MW (6946.01).

Insulin (M_MONO_ 5729.6009; M_av_ 5734.51) and apomyoglobin (M_MONO_ 16940.9650; M_av_ 16951.49) were used as internal standards for mass calibration while acquiring spectra. The M_av_ of Hpn was determined as 6945.66±0.34 Da calculated from the average of at least five different measurements. A representative calibrated peak of Hpn[M^+^] with *m/z* 6946.05 is shown in **Fig 3B**. However, MALDI-TOF-MS analysis confirmed that the M_av_ of purified recombinant Hpn was 6945.66±0.34 Da instead of the calculated molecular mass of 7,077 Da. This suggests the loss of the N-terminal methionine residue (*m/z* 149.21), probably in post-translation process in *E. coli* and this observation is consistent with the previous reports [4,5].

### Hpn exhibits both “gel shifting” and “metal gel-shift”

The reduction by β-mercaptoethanol and boiling before gel loading of purified Hpn had no effect on the appearance of two bands even in different polyacrylamide-gel concentrations [(~14 kDa and ~18 kDa on 15% SDS-PAGE; **Fig 1C-E**) and (~7 kDa and <7 kDa on 20% SDS-PAGE; **Fig 4A**)]. This was further analyzed by Ni^2+^ addition and removal (using EDTA) separately to the purified Hpn. In both cases, samples with and without boiling were analyzed in order to assess if Hpn showed a characteristic property of “heat modifiability” like outer membrane proteins (OMPs) from *E. coli* [51]. The concept of heat modifiability constitutes preservation of both folded and unfolded structures upon SDS treatment to purified OMPs but with no heat and shows two different bands on SDS-PAGE. Heating denatures the folded β-content of OMPs and SDS-PAGE data shows single band. No difference in migration pattern was observed between heated and non-heated samples of Hpn indicating the absence of heat modifiability (**Fig 4A**).

**Fig 4.**
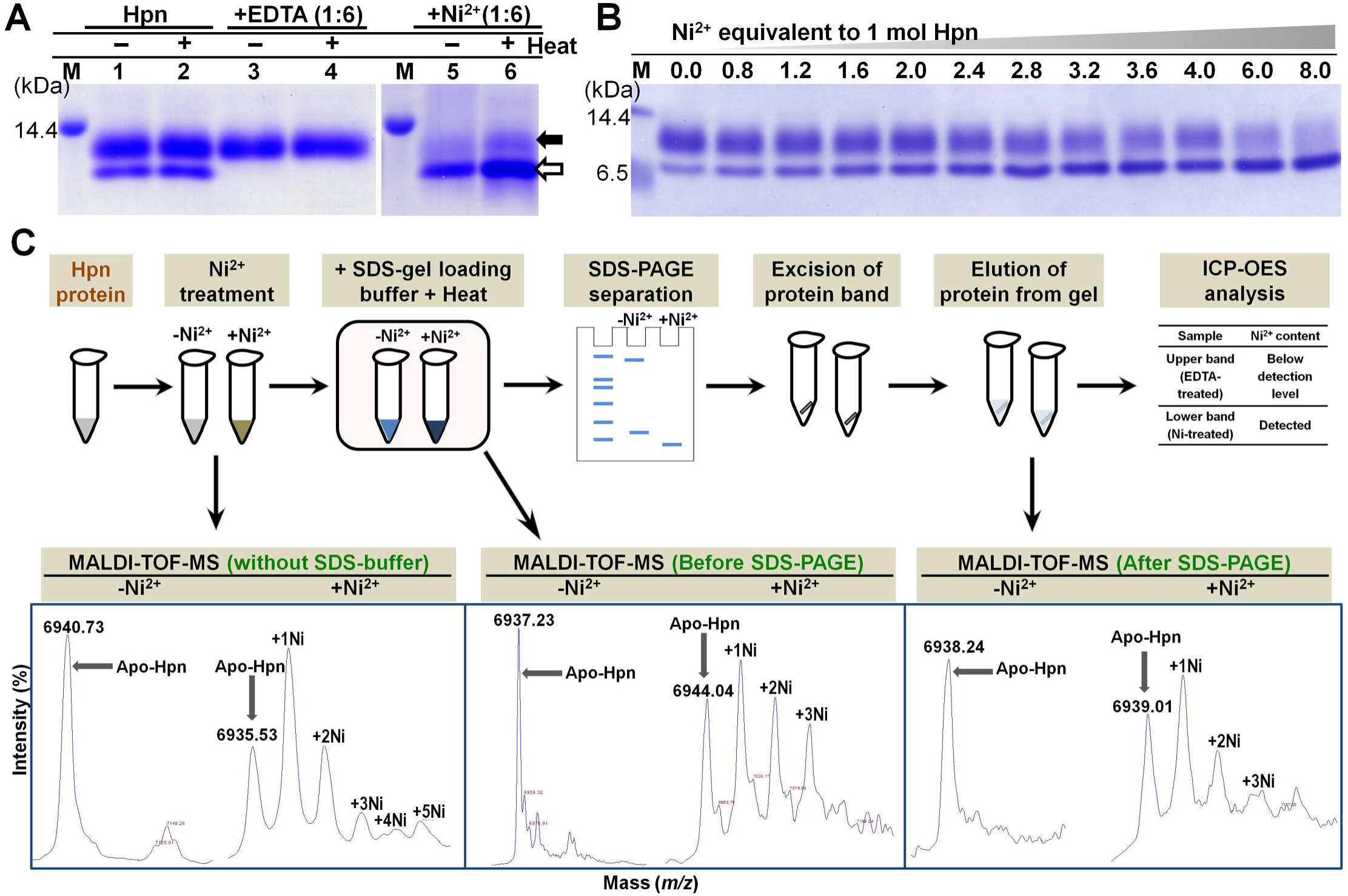
Confirmation of “Metal gel-shift” mechanism. A. Effect of EDTA and Ni^2+^ ion treatment on migration rate of recombinant Hpn in SDS-PAGE (polyacrylamide-gel 20%). Lane M, protein marker; lane 1 and 2, Hpn before and after boiling (3 min at 100ºC), respectively; lanes 3 and 4, EDTA-treated Hpn without and with boiling, respectively; lanes 5 and 6, Ni^2+^-treated Hpn without and with boiling, respectively. B. The SDS-PAGE analysis of partially-metalated-Hpn (25 µM) treated with increasing concentration of Ni^2+^ (1:0, 1:0.8, 1:1.2, 1:1.6, 1:2.0, 1:2.4, 1:2.8, 1:3.2, 1:3.6, 1:4.0, 1:6.0 and 1:8.0). Equal volume of heat-denatured protein applied in each lane. C. Scheme used for MALDI-TOF-MS analysis of Hpn protein that was heat denatured in Laemmli buffer. MS data was measured for Hpn treated with or without Ni^2+^ ion (1:6 mol equivalent ratios). Further, MS data for Hpn (with or without Ni^2+^) treated in Laemmli buffer (before and after SDS-PAGE) was measured. Even though some interference due to adducts was observed in samples treated with Laemmli buffer or gel-eluted fractions, metalated peaks (showing Hpn-Ni^2+^ complexes) were distinct. The occurrence of metalated peaks was observed only for Ni^2+^-treated Hpn in all the conditions.

Moreover, EDTA-treated Hpn showed only one band of ~7 kDa on 20% polyacrylamide-gel. On the other hand, the only protein band observed in Ni^2+^-treated Hpn was of compact lower-size <7 kDa. Appearance of only one of the either band demonstrates that both bands were derived from same protein i.e. Hpn. The partially-metalated Hpn treated with relatively lower mol equivalent proportion of Ni^2+^ (0.0, 0.8, 1.2, 1.6, 2.0, 2.4, 2.8, 3.2, 3.6, 4.0, 6.0 and 8.0) demonstrated gradual shift with decrease in homogeneity of upper band followed by increased intensity of compact lower band (**Fig4B**).

The Hpn protein treated for SDS-PAGE analyses was directly used for MALDI-TOF-MS measurements and mass spectra of Ni^2+^-treated Hpn only showed peaks for protein-metal complexes (**Fig 4C**). The mass spectra of protein fractions eluted from SDS-gel corresponding to ~7 kDa and compact <7 kDa on 20% gel were measured. Only lower compact band (<7 kDa) confirmed mass spectra for partially preserved protein-metal complexes (**Fig 4C**). This MS data prompted us to quantify the metal-content of SDS-gel-eluted bands. The Ni^2+^ content in lower compact bands (combining several gel slices together) was detectable in ICP-OES. Measurements for EDTA-treated Hpn band were below the detection limit signifying no metal bound to Hpn protein.

Several studies have reported preservation of protein-metal complex in SDS-PAGE **(supporting information, S2 Table in S1 file)** and also in MALDI technique (**Table 3**). Thus, those studies have provided experimental evidence that some metal-binding proteins/metalloproteins can retain protein-metal complex even after using “harsh” experimental protocols (i.e. use of SDS, denaturing agents, acidic matrices, organic solvents or heating). Therefore, it is postulated that the methodology employed in the present study was able to detect Hpn-Ni^2+^ complex and it was not a consequence of artifact formation. Previous circular dichroism (CD) studies have reported more compact structure (an increase in β-sheet with reduced α-helical content) after Ni^2+^ binding to Hpn [5,6].

**Table 3.**
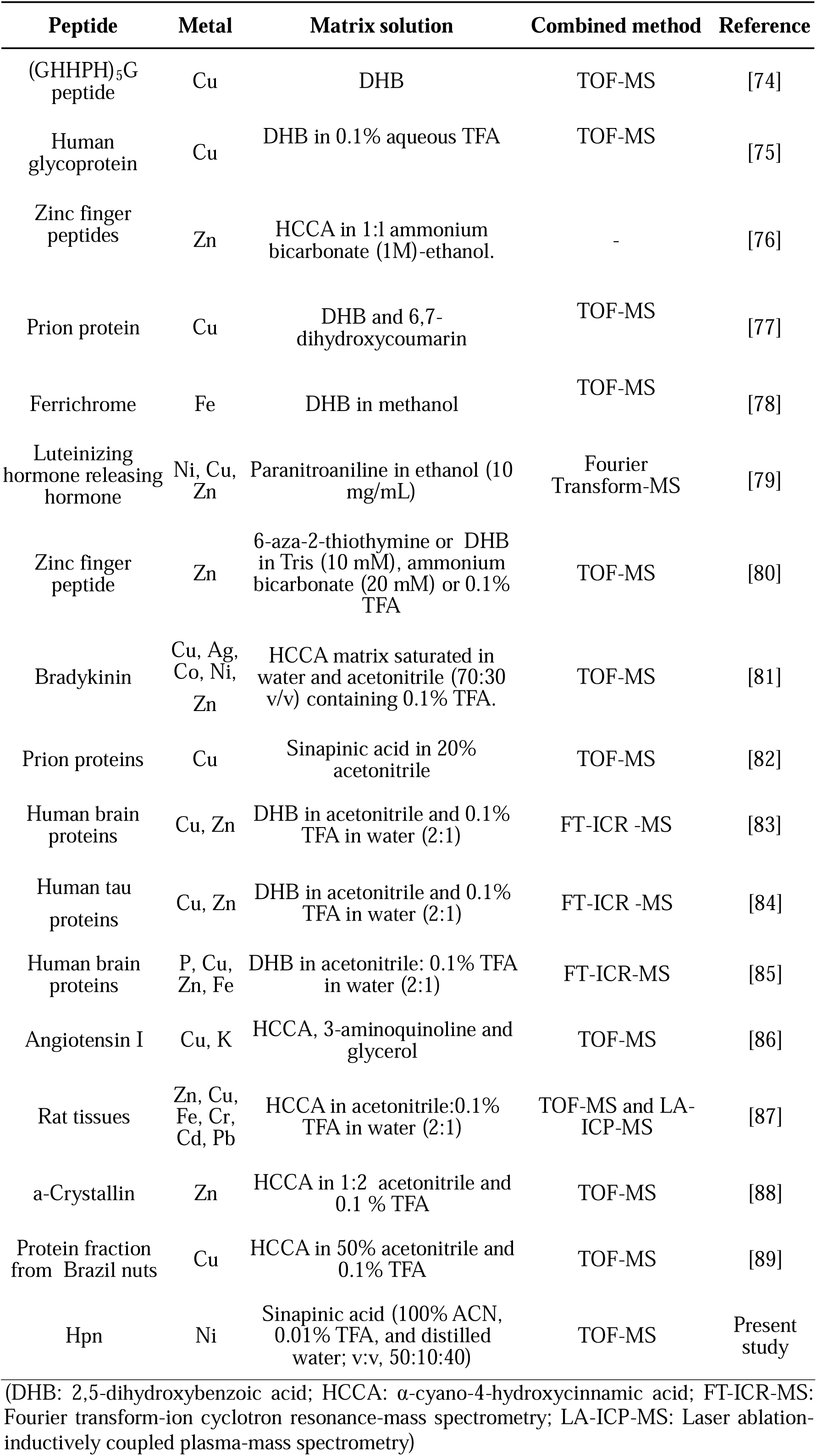
List of non-covalent protein-metal ion interactions studied by MALDI in combination with other methods.

Taken together, these data indicate that preserved protein-metal complex forms a more compact structure leading to faster migration on SDS-PAGE. This phenomenon of reversible shift in position when treated with either EDTA or Ni^2+^ was termed as “metal gel-shift”.

### Polyacrylamide-gel concentration determines migration rate of Hpn on SDS-PAGE regardless of “metal gel-shift”

Polyacrylamide-gel concentration can dictate the migration speed of some polypeptides [31]. To test the hypothesis that “metal gel-shift” is related to only Ni^2+^ binding to Hpn or whether gel percentage has an effect on it, we thoroughly analyzed gel mobility of apo-Hpn and Ni^2+^-treated Hpn with different polyacrylamide-gel concentrations. This concept was investigated with purified Hpn after removing trace amounts of Ni^2+^ (**Fig 5A**). The migration distance of apo-Hpn and Ni^2+^-treated Hpn relative to marker proteins (Nacalai Tesque, Inc. Japan) was analyzed using ImageJ software [52] (**protocol provided in Supporting information, Annexure A in S1 file**). Outcomes from these experiments were in agreement with our hypothesis; the Ni^2+^-treated Hpn migrated faster than untreated Hpn showing MW difference of 3-4 kDa (**Fig 5B**). These results demonstrated that apo-Hpn migrated slowly in contrast to Ni^2+^-treated Hpn in all analyzed polyacrylamide-gel concentrations, signifying that “metal gel-shift” is intimately associated with Ni^2+^-binding to Hpn. This also shows that higher polyacrylamide-gel concentrations resulted in faster migration of both the forms on SDS-PAGE (with and without Ni^2+^).

**Fig 5.**
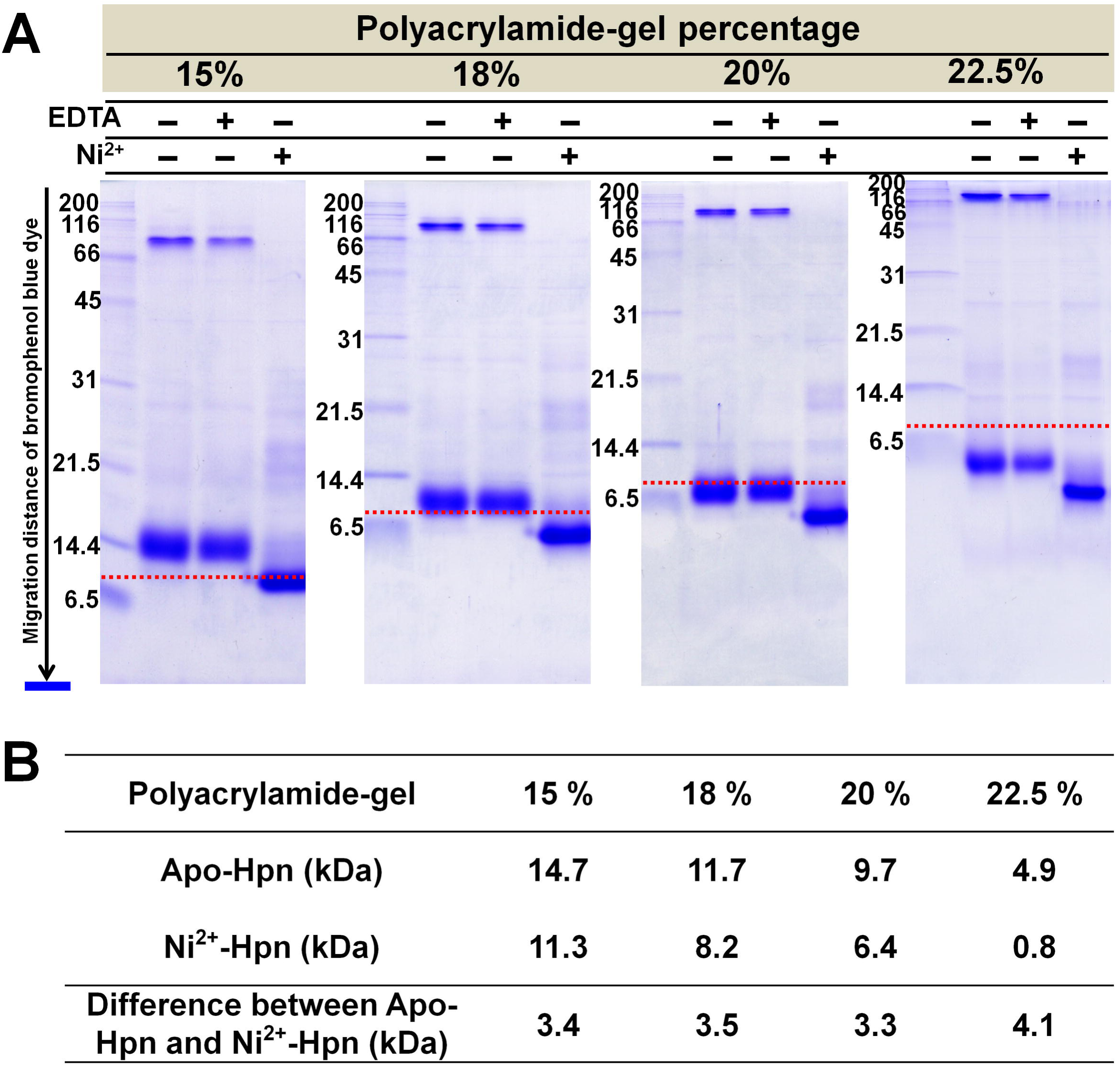
Effect of polyacrylamide-gel concentration on migration speed of recombinant Hpn without or with metal-treatment on SDS-PAGE. A. Relative position of apo- and Ni^2+^-treated Hpn protein depending on the polyacrylamide percentage (15, 18, 20 and 22.5%) in gels. The SDS-gel electrophoresis was done till bromophenol blue dye reached to the bottom in all gels. Therefore, theoretical MW values for marker proteins-lysozyme (14.4 kDa) and Trypsin inhibitor (21.5 kDa) were taken into consideration for estimation of apparent MW using ImageJ software (protocol provided in Supplementary information as Annexure A). Red dotted line depicts expected position corresponding to theoretical MW (~6.9 kDa). B. The apparent MW of apo- and Ni^2+^-treated Hpn separated on different polyacrylamide-gel concentrations was estimated by comparing relative migration distance of Hpn with globular marker proteins on SDS-PAGE.

### Analysis of non-covalent Hpn-Ni^2+^ complexes

We evaluated inter-conversion of Hpn-metal complexes using purified Hpn by MALDI-TOF-MS. Although non-covalent complexes are expected to be stable at physiological pH, acidic conditions or organic solvents used in matrix preparation may not be the only factors important in the dissociation and prevent detection of these complexes in MALDI analysis [53]. In this work, we applied full-length Hpn protein for studying Ni^2+^ binding. First, we standardized the set of conditions for acquisition of spectra for intact Hpn, and then we investigated different types of matrices to analyze protein-metal ion complexes. Among the tested combinations of acidic and non-acidic matrices, we found a mildly acidic matrix (100% ACN, 0.01% TFA, and distilled water; v:v, 50:10:40) was most suitable in our experimental conditions, and this was used in subsequent spectral measurements. This is the first study reporting the successful application of MALDI-TOF-MS for investigating protein-Ni^2+^ ion complexes. The MS data showed the progressive appearance of all possible metal-bound species of Hpn, signifying that binding of metal ions at each site on protein may occur independently. Moreover, a decrease in spectral intensity of the apo-protein with increasing amounts of Ni^2+^ and increased intensity of metalated species provided conclusive evidence for the binding of Ni^2+^ to Hpn.

To evaluate Ni^2+^ binding, Hpn treated with increasing molar concentrations of NiSO_4_ solution was analyzed. Generally, the peaks of protein-metal complexes are smaller than the peak of the apo-protein, even if a protein-metal interaction is already known to exist. Therefore, higher amounts of Ni^2+^ were added to Hpn after confirming that no obstruction was caused by it in the MALDI spectrum. Ni^2+^ solution was added to apo-Hpn (25 µM), pH 7.5 at an increasing mol equivalent ratio of Hpn to Ni^2+^ ion. After 1 h incubation at room temperature, protein-metal solution was mixed with matrix and MS were acquired as described above (**Fig 6**).

**Fig 6.**
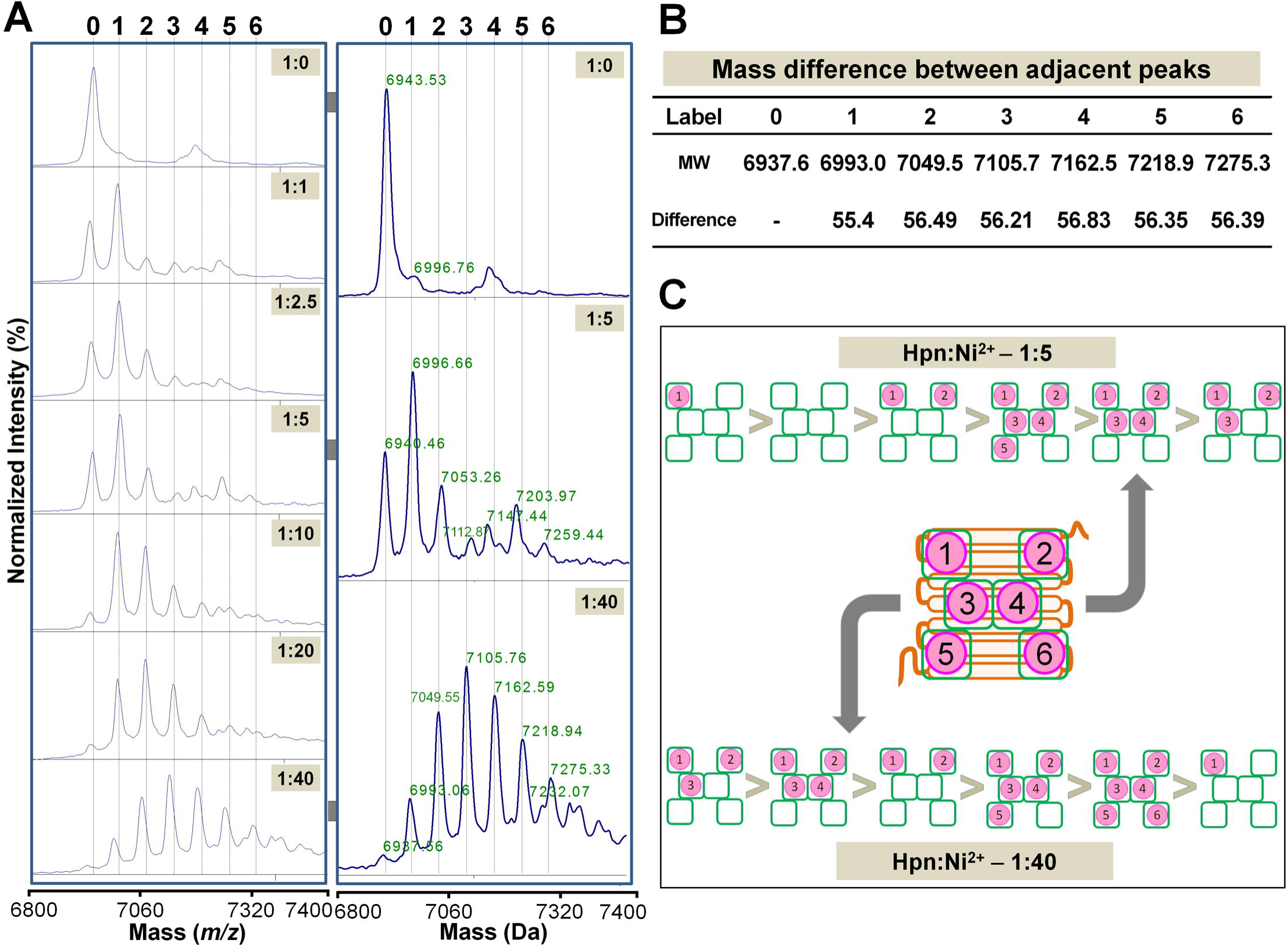
MALDI-TOF-MS analysis of Ni^2+^ binding to Hpn. A. Spectra obtained with increasing concentrations of Ni^2+^ (1:0, 1:1, 1:2.5, 1:5, 1:10, 1:20 and 1:40, mol equivalent from top to bottom, respectively) added to apo-Hpn (25 µM) shown in the right panel, and enlarged view of three representative spectra (1:0, 1:10 and 1:40) shown with molecular mass of each peak in the left side. Numbers above dotted line correspond to the number of Ni^2+^ bound to Hpn protein. B. Mass difference calculated between adjacent peaks was found to be approximately equal to the molecular mass of the Ni^2+^ ion (58.69) with the loss of two H^+^ atoms (molecular weight [MW] of H^+^ =1.00794) upon metal binding. C. Model of Ni^2+^ ion binding to Hpn. Order of occurrence has drawn on the basis of peak intensity obtained in MALDI-TOF-MS data.

Representative MS of metalated species of Hpn clearly detected a progression with seven major peaks (at *m/z* 6937.66, 6993.06, 7049.55, 7105.76, 7162.59, 7218.94, and 7275.33). The MS showed peaks of diverse intensity ranging from two to six Ni^2+^ ions bound to Hpn (**Fig 6A**), with no specific species dominant in all the measured protein to metal ratios. The mass differences between adjacent peaks were about *m/z* 56.7 (**Fig 6B**), matching the added Ni^2+^ ion (*m/z* 58.69 of Ni^2+^) with the loss of two hydrogen residues during the ionization (*m/z* 1.007825×2). Overlay analysis of MS with and without Ni^2+^ showed slight mass differences (*m/z*) between expected and observed molecular masses may be due to tightly bound ions as previously observed for UreE protein from *H. pylori* [54]. The seventh and subsequent peaks were bifurcated possibly because of matrix adducts interference and hence considered not reliable for assignment to Hpn-Ni^2+^ ion complex. Titration of Hpn with relatively lower Ni^2+^ concentrations (1:5) indicated at least one preferential higher-affinity site, peak intensities of the protein-metal complexes were not according to the loading number of Ni^2+^ ions on binding sites of Hpn (1>0>2>5>4>3). Furthermore, a difference in peak intensity with higher amounts of Ni^2+^ (1:40) was observed (3>4>2>5>6>1) suggesting the presence of several binding sites with different affinity. The progressive appearance of peaks for all of the possible Hpn-Ni^2+^ ion complexes implied that Ni^2+^ binding to Hpn occurred in a non-cooperative way (**Fig 6C**) as reported in cysteine-rich human metallothionein 1a [55] and histidine-rich *E. coli* SlyD protein [21].

### Ni^2+^ tolerance and accumulation in Hpn-expressing *E. coli* cells

If there is higher metal accumulation inside cells expressing the recombinant metal-binding protein, then it should show better cell growth with higher tolerance compare with wild-type upon addition of higher amounts of metal to the culture. However, this phenomenon was not observed in the current as well as previous studies in Hpn-producing *E. coli* cells (Hpn+) compared with wild-type (Hpn–) grown in LB medium [5]. Even though Hpn have more efficient metal-binding ability as observed in MALDI-TOF-MS analysis, higher Ni^2+^ accumulation measured by ICP-OES (6× for Hpn+ compared with Hpn–) was not correlated with higher tolerance and cell survival (<twice in Hpn+ compared with Hpn–) (**Fig 7A-C**). We hypothesized that an unknown composition of nutrients in LB medium may compensate for toxicity or availability of free Ni^2+^ ion to cells. Therefore, we tested M9 medium with a defined nutrient composition. As shown in **Fig 7D-E**, there was almost no further growth of wild-type (Hpn–) at 50 µM and 100 µM Ni^2+^ supplied in M9 medium but Hpn producing cells (Hpn+) were grown normally with approximately 30 and 45 times higher intracellular Ni^2+^ content respectively (**Fig 7F**).

**Fig 7.**
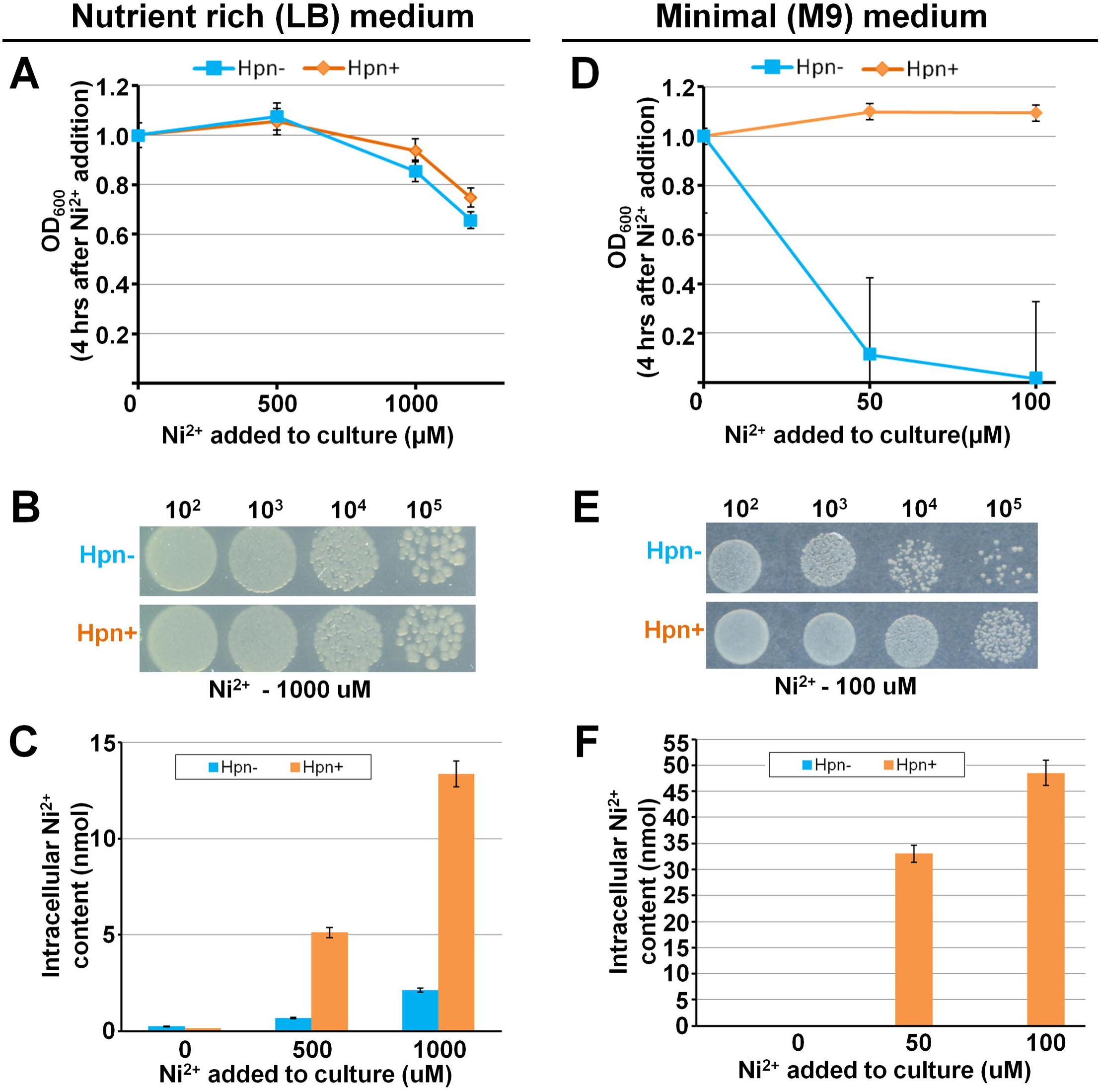
Ni^2+^ tolerance, cell survival, and accumulation in Hpn-expressing *E. coli*. Effect of Ni^2+^ on growth of *E. coli* in Luria-Bertani (LB) (panel A) and M9 medium (panel D). Growth curve was plotted in terms of optical density (OD) against amount of Ni^2+^added to culture. Cell survival under Ni^2+^ stress analyzed by dots blot as shown in panel B (LB) and E (M9). Panel C (LB) and F (M9) represent the intracellular Ni^2+^ content in *E. coli* expressing *hpn* gene compare to that of without *hpn* gene.

### Discussion and Conclusions

The main findings of current work are as follows: 1) Hpn forms multimers in native state as well as SDS-resistant multimer when eluted with 400 mM imidazole in buffer; 2) Ni^2+^-binding to Hpn altered antibody binding in western blot and ELISA signifying differential exposure of His-rich region upon metal-binding; 3) higher polyacrylamide-gel concentrations resulted in faster migration of Hpn in SDS-PAGE; 4) average molecular mass of Hpn determined as *m/z* 6945.66±0.34; 5) Hpn forms SDS-resistant (preserved) protein-metal complexes and exhibits metal-triggered shift in electrophoretic mobility causing “metal gel-shift” mechanism; 6) MALDI-TOF-MS was effectively employed to study non-covalent Hpn-Ni^2+^ ion complexes showing up to six Ni^2+^ ions bound per monomer in a non-cooperative way suggesting an equilibrium between Hpn-metalated species dependent on metal availability. These findings explore various unusual physicochemical aspects of Hpn. Also, higher tolerance and Ni^2+^ accumulation in Hpn-expressing *Escherichia coli* than wild-type in minimal (M9) and nutrient-rich (LB) supply suggests protective role and potential to load higher amounts of Ni^2+^ that corroborating with MALDI-TOF-MS data.

The Hpn protein contains remarkably high number of histidine residues, mostly in clusters (**Fig 1A**). Histidine is unique in its molecular structure with an imidazole ring in its side chain (an aromatic motif), and this can act as a ligand for metallic cations and as a hydrogen bond donor or acceptor. “Stacking” behavior of the aromatic rings of histidine can be one possible mechanism responsible for the formation of non-covalent multimeric complexes; however, this concept is yet to be understood clearly in multimeric proteins [56]. Elution buffer containing imidazole (400 µM) led to formation of SDS-resistant multimer that survived even in reduction and heat denaturation but converted to apparent monomer in denatured Ni^2+^-treated Hpn (**Fig 5**). The preserved SDS-resistant protein-protein oligomer is reported for several proteins under certain chemical environment (**Supporting information, S1 Table in S1 file**) [57–62].

### Gel shifting

Protein migration can be affected by several factors including molecular size, shape, net charge, MW of the protein, and polyacrylamide concentration [31]. SDS normally binds at hydrophobic sites, therefore it is reasonable that denatured apo-Hpn migrates at a slower rate compared with marker proteins, because of the higher amount of hydrophilic residues and only one hydrophobic amino acid in the Hpn [23,25,28]. A smaller protein with a higher number of hydrophilic residues may have a greater hydrodynamic radius than a larger protein but weaker hydration [63]. Similar results are reported for cystic fibrosis transmembrane conductance regulator for the reason that of a change in helical structure altered SDS-binding indicating protein-SDS complex size was a more important factor than net charge [23] and its interaction with the sieving effects of a polyacrylamide gel [25].

The SDS-PAGE data demonstrated that the polyacrylamide-gel concentration affects the migration rate of Hpn. Similar results are observed in “gel shifting” patterns for transmembrane proteins in SDS-PAGE [31]. This indicate as any factor that changes effective molecular size and net charge of protein-SDS complex can affect migration speed depending on polyacrylamide-gel concentration, specifically affecting its interaction with the available space in SDS-gel matrix [23,24,31]. In case of Hpn, impact of “stacking” behavior seems to exceed all other factors and may control the migration speed depending on polyacrylamide-gel concentration. Taken together, interplay between protein-protein interaction and sieving effects of gel matrix may be a significant factor that influences gel mobility.

### Metal gel-shift

Generally, non-covalent interactions should disrupt during SDS-PAGE owing to activity of SDS and reducing agent [60]. However, preservation of protein-metal complex in SDS-PAGE is reported in several cases, possibly due to incomplete or “reconstructive denaturation” (**Supporting information, S2 Table in S1 file**) [35,36,60,62,64–69]. Preservation of partial Hpn-Ni^2+^ complex even after electrophoretic separation implies significant strength and stability of Hpn-Ni^2+^ bond. Although exact mechanism of resistance to reduction or denaturation and retaining metal ion is not yet known, similar results were observed in case of platinum-binding proteins [70].

The change in electrophoretic mobility on SDS-PAGE after metal-binding to a protein is rarely acknowledged except for some Ca^2+^-binding proteins including calmodulin isoforms [36,37]. The wild-type and recombinant CDPKs from soybean [38], tobacco [39] and Arabidopsis [40] display shift in electrophoretic mobility on SDS-PAGE and migrate faster or slower depending on Ca^2+^ availability and is probably due to Ca^2+^-induced conformational change [40]. The protein band of apo-Hpn was not as sharp as that of Ni^2+^-treated Hpn, which might be caused by conformational changes upon metal binding. Thus, higher degree of compactness or somehow altered SDS-binding to protein-metal complex that may allow faster migration of Ni^2+^-treated Hpn than apo-Hpn. Faster migration on SDS-PAGE of unreduced against reduced lysozyme [71] and for a membrane protein OmpA [72] also suggested a possible role of structural compactness in “gel shifting”. Considering added MW of metal ions, theoretical MW of Hpn protein in “metal-gel shift” band is higher than apo-Hpn band. On the contrary, “metal-gel shift” band is migrating faster to lower position compared to apo-Hpn band indicating conformational change (and not actual MW) is decisive in governing migration rate.

From the experimental results, mechanism of “gel shifting” and “metal gel-shift” can be described as shown in **Fig 8**. Anomalous migration of apo-Hpn (scheme highlighted with yellow background) or metal-bound Hpn (scheme highlighted with green background) might be due to differential amounts of SDS bound to protein, histidine stacking, altered hydrodynamic radius, degree of compactness after metal-binding, or a combination of either of these factors. The different structures are illustrated and discussed thoroughly in **Fig 8**. Nevertheless, there might be additional biochemical and biophysical processes governing differential protein migration in SDS-PAGE that is not explained in this scheme.

**Fig 8.**
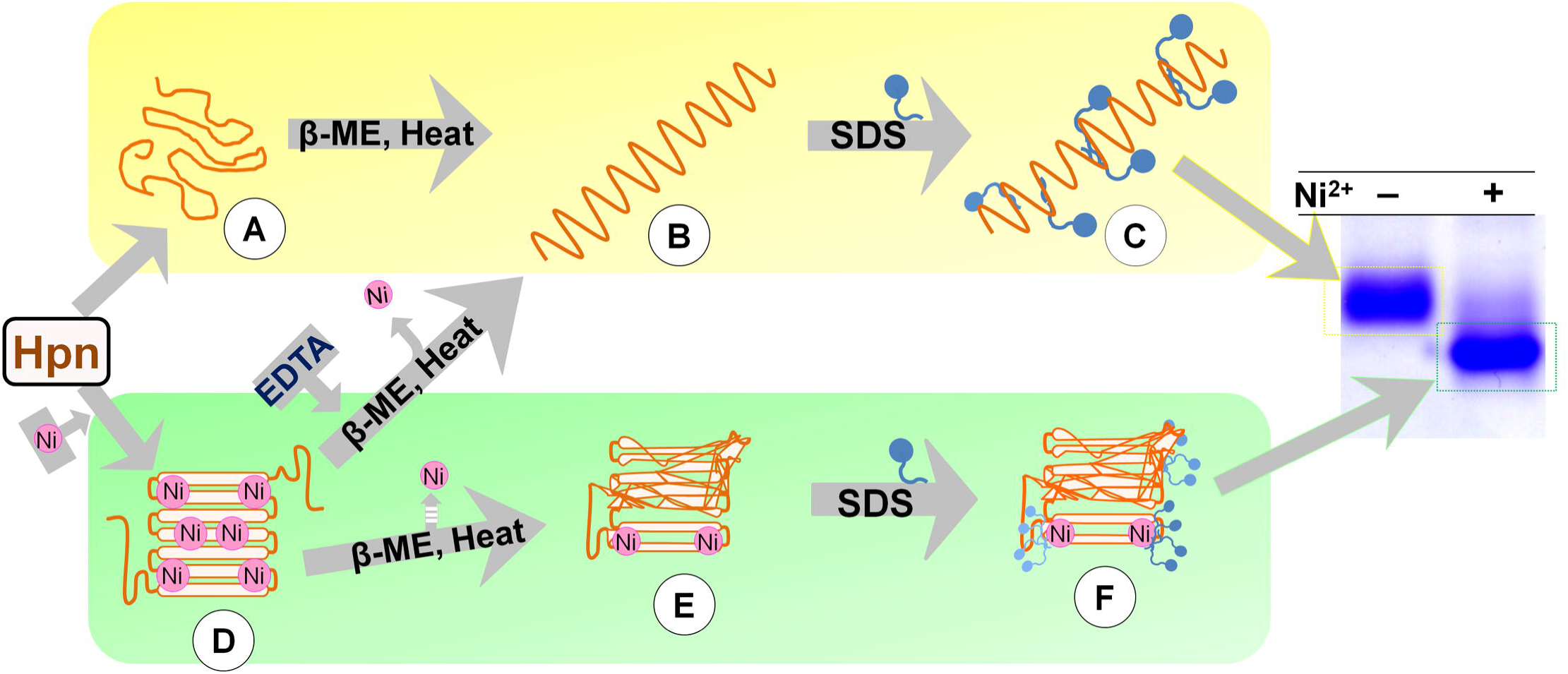
Probable interrelationship between differential electrophoretic mobility of Hpn and Ni^2+^ binding. Hpn may not have a definite form in the absence of Ni^2+^ (A). After denaturation (B), smaller amounts of SDS binding/stacking behavior/larger hydrodynamic radius as well as a combination of some or all of these conditions (C) might have resulted in slower migration on SDS-PAGE (scheme highlighted with yellow background). Ni^2+^-treated Hpn forms a more compact structure (D). Pictorial structure of metalated Hpn is drawn to explain the model. MALDI spectra showed a partial Ni^2+^ bound form (E) in denatured SDS-PAGE. Altered binding of SDS (F) caused by replacement of protein-protein to protein-SDS contacts (inhibiting stacking behavior) and/or degree of compactness (or reduced hydrodynamic radius) may be key factors responsible for “metal gel-shift” (scheme highlighted with green background). β-ME, β-mercaptoethanol; EDTA, ethylene diaminetetraacetic acid.

The binding of non-denatured Hpn to Ni^2+^ column and reversible “metal-gel shift” in SDS-PAGE presented herein together with previous CD studies [5] indicated that Hpn protein could easily exchange metal ion suggesting the position of metal-binding domains are solvent exposed and present at Hpn surface. Metal-induced structural changes leading to the formation of a definite form has been reported for several other proteins, mostly associated with β-turns [21,73]. Accordingly, metal-binding region (not entirely hidden inside the structure of protein) could easily accessible for metal exchange and may involve small but unique structural rearrangements.

### Metal gel-shift and MS data

The pre-requisite for successful application of any mass spectrometry technique while studying protein-metal interactions is the optimization of suitable method that preserves non-covalent complexes. The MALDI technique has been used previously to study non-covalent protein complexes (**Table 3**) by adjusting the range of parameters in order to preserve the non-covalent interactions during acquisition of spectra [46,47,90].

Using mild-acidic matrix, spectral measurements showed distinct peaks, validating one apo-Hpn peak with six of Hpn-Ni^2+^ complexes. Singly charged species of Hpn in MALDI-TOF-MS might allow detection of a sixth Hpn-Ni^2+^ complex in mild-acidic conditions that not reported in previous studies [6,7]. However, appearance of weaker or non-specific protein-metal complexes due to higher amounts of metal added to protein samples is a limitation of the ionization process in positive mode measurements [21] and this possibility for sixth Hpn-Ni^2+^ complex cannot be excluded. The interfaces of metal ions with histidine-rich peptides have not been investigated so far, perhaps because such peptides are not possible to study with routinely used techniques owing to overlapping signals [91]. Even though MS data alone are not sufficient to interpret the exact mechanism of metal-binding to various sites, these data have revealed the high flexibility (plasticity) of Hpn for Ni^2+^ binding and its potential to load higher amounts of Ni^2+^ in such a small structure of ~7 kDa.

Our MS data imply several key points that are complementary to “metal gel-shift”. At first, all available Hpn molecules were in metalated form at higher Ni^2+^ in MS data. Similarly, presence of only “metal-gel shift” band at higher Ni^2+^ was observed on SDS-PAGE (**Fig 4B**). Second, Hpn showed MS peaks for metalated species even treated with lower metal concentrations (**Fig 6**). This coincides with the appearance of "metal-gel shift" band in all protein-metal molar ratios (lower to higher) on SDS-PAGE (**Fig 4B and C**). Third, mass spectral intensity for apo-Hpn gradually decreased in Ni^2+^-treated Hpn and subsequently mass spectral intensity for either of the metalated species was increased in respective measurements showing progressive appearance of Hpn-Ni^2+^ peaks. This is also consistent with the gradual increase in heterogeneity of apo-Hpn band followed by compact "metal-gel shift" band.

Interestingly, at relatively higher amount of Ni^2+^, distribution of mass for each Hpn-Ni^2+^ complex was highly heterogeneous in MS data but on contrary, SDS-PAGE showed only one compact protein band. Electrophoretic separation of metalated species having smaller mass difference of each added Ni^2+^ (0.055 kDa) cannot be distinguished on regular SDS-PAGE owing to limited resolution. Also, several factors can affect the migration of each metalated species of Hpn on SDS-PAGE including consequence of boiling, Laemmli buffer components, re-arrangement of protein-protein, protein-metal, and/or protein-SDS interactions. But this is not the case in MS measurements. Moreover, previous studies suggested involvement of almost all the four cysteine residues in Ni^2^-binding [5,10–12]. Therefore, partial or complete reduction of disulfide bonds in two pairs of cysteine residues may release some but not all metal ions bound to Hpn during “constructive” denaturation. It may produce partially-metalated Hpn species in equilibrium (observed in MS of protein fractions before and after SDS-PAGE, **Fig 4C**) that migrated as homogenous “metal gel-shift” band on SDS-PAGE. In addition, differential ionization efficiency and ion suppression effect in MS may interfere in mass-to-charge ratio [92]. The thermodynamic structures of the apo-protein for metalation and the fully metalated protein for demetalation may not be the same [21]. Thus, without further structural studies, it is intricate to compare precise mass distribution of metalated species on SDS-PAGE and further studies are under investigation. Nevertheless, “metal-gel shift” together with MALDI-TOF-MS data establishes that the positional shift is directly associated with metal-binding to Hpn and it is reversible upon metal removal.

In brief, our study reveals a novel mechanism of “metal gel-shift” responsible for shifts in electrophoretic gel mobility of Ni^2+^-treated Hpn on SDS-PAGE signifying metal-induced conformational changes. This property can be used to explore interactions between histidine-rich proteins and surfactant to investigate how metal-binding to a histidine-rich protein changes its confirmation and hydrophobic or electrostatic interactions.

## Conflict of interest

The authors declare that they have no conflicts of interest with the contents of this article.

## Author contributions

HH and EHM conceived the idea for the project, supervised the study, provided essential reagents and edited the manuscript. RMS conducted most of the experiments, analyzed the results and wrote the manuscript. YI and JM conducted *H. pylori* culture experiments including genome DNA extraction and gene cloning. All authors edited the manuscript, reviewed the results and approved the final draft of this manuscript.

## Acknowledgement

The protocol and facility to perform ELISA was kindly provided by Dr. T. Tsuboi and Dr. E. Takashima (Division of Malaria Research, Proteo-Science Center, Ehime University, Matsuyama, Ehime, Japan).

## S1 Supporting Information

Table S1. Proteins showing apparent SDS-resistant oligomeric forms upon denaturing SDS-PAGE.

Table S2. List of proteins retaining metal ion on SDS-PAGE.

Fig S1. Schematic diagrams of gene constructs and protein analysis by SDS-PAGE used for western blot and ELISA analysis.

A. Schematic diagram of gene construct consisted of *gfp* fused with artificial His.tag (left panel) and GFP-His6 protein expression confirmed by SDS-PAGE (right panel).

B. Schematic diagram of *gfp* fused with *hpn* at C-terminal followed by stop codon (left panel) and recombinant protein of GFP-Hpn fusion analyzed by SDS-PAGE (right panel).

Fig S2. Comparison of DNA sequence of *hpn* from *Helicobacter pylori* strain SS1 (this study) with strain 26695 (NCBI data).

The promoter region and *hpn* gene of strain SS1 was PCR amplified and nucleotide sequence was compared with NCBI data of strain 26695 (GenBank accession number U26361). Putative promoter elements are shown in a box. The *hpn* gene region is shown in uppercase letters and mutations at the nucleotide level are shaded in gray. The putative terminator region of transcription is underlined. The promoter region was highly conserved in strain SS1 including Shine-Dalgarno sequence (GGAG) and promoter elements (-10 and -35).

Fig S3. Separation of purified Hpn on non-denaturing blue native-PAGE.

High molecular weight marker (GE Healthcare), abbreviated as HMW, was used in all gels [Thyroglobulin (669 kDa), Ferritin (440 kDa), Catalase (232 kDa), Lactate dehydrogenase: (140 kDa) and Albumin (66 kDa)]. Recombinant Hpn (200 µM) was treated with either 1 mM of EDTA or Ni^2+^ independently and then applied to 10% native-gel. Apparent multimeric complexes of >670, ~500, ~230 kDa were observed in presence or absence of Ni^2+^ and EDTA.

## Annexure S1

Protocol followed for analysis of apparent MW of Hpn on SDS-PAGE in Fig. 5.

